# Practical utility of sequence-to-omics models for improving the reproducibility of genetic fine-mapping

**DOI:** 10.64898/2026.02.04.703796

**Authors:** Michael D. Sweeney, Hyun Min Kang

## Abstract

Recent advances in deep learning have led to the development of sequence-to-omics (S2O) models that predict molecular phenotypes directly from DNA sequences. Here, we systematically evaluate the utility of these models, e.g., AlphaGenome, Borzoi, Enformer, and Sei, for improving the reproducibility of genetic fine-mapping across expression quantitative trait loci (eQTL) datasets from Genotype-Tissue Expression (GTEx), Trans-Omics Precision Medicine (TOPMed), and Multi-Ancestry Analysis of Gene Expression (MAGE) projects. We show that purely statistical fine-mapping often yields high replication failure rates (RFRs), but integrating S2O model predictions substantially reduces RFRs and enhances the accuracy of prioritizing SNPs replicated in other consortia. We describe a generalized framework for functionally informed fine-mapping that combines traditional posterior inclusion probabilities (PIPs) from statistical fine-mapping methods with scores from S2O models to generate functionally informed PIPs (fiPIPs) that improve reproducibility. Our findings demonstrate that S2O models, particularly newer ones like AlphaGenome and Borzoi, enable robust identification of replicated variants across consortia, highlighting their promise for scalable, functionally aware genetic mapping.

## Introduction

Recent advancements in deep learning methods in genomics have introduced powerful new tools for predicting the functional impact of genetic variation on molecular and complex traits, orthogonally to genetic association studies. For example, protein language models, such as AlphaMissense^1^, Evolutionary Scale Modeling (ESM)^2^, and others^3–11^, have achieved higher accuracy in identifying protein-disrupting and pathogenic missense variants than methods that have been the standard for decades^12–14^. Trained on millions of protein sequences using Transformer-based architectures, these models learn the evolutionary constraints and structural relationships embedded within proteins. This allows them to accurately assess how specific amino acid substitutions affect protein stability and function. Similarly, genomic language models like Evo 2^15^, GPN-MSA^16^, HyenaDNA^17^, DNABERT-2^18^ and others^19–22^, use cross-species DNA language models^23^ to model the language of the entire genome to extend these functional impact predictions beyond protein-coding variants.

A distinct but related class of models, sequence-to-omics (S2O) models like AlphaGenome^24^, Borzoi^25^, Enformer^26^, and Sei^27^, and others^28–30^ are trained jointly on DNA sequences and multiple epigenomic profiles^31,32^. By using convolutional neural networks, transformers, or state space models, they can predict tissue-specific, multi-omic outcomes—such as transcriptional levels, chromatin accessibility, and histone modifications—from any given DNA sequence, also facilitating prediction of the molecular impact of arbitrary genetic variants, including common, rare, or even previously unobserved variants. However, it is important to note that multiple studies report that some of these models are not yet as accurate as statistical models from population-scale transcriptome-wide association studies (TWAS) for predicting personal gene expression levels^33,34^.

In this paper, we focus on the practical utility of these emerging S2O models for a critical challenge in human genetics: improving the reproducibility of genetic fine-mapping. Statistical methods like SuSiE^35^ and FINEMAP^36^ are widely used to pinpoint putative causal variants from genetic association signals. However, these methods often suffer from much lower reproducibility between independent studies than their statistical confidence scores would suggest^37^. This inconsistency, likely driven by factors such as the imperfect modeling of causal variant effect distributions, linkage disequilibrium, imputation uncertainty, and rare causal variants, hampers the reliability of downstream analyses like polygenic risk score construction^38^, target gene prioritization^39,40^, Mendelian Randomization^41^ and colocalization^42^.

While previous work has shown that incorporating functional predictions can improve the statistical power of fine-mapping studies^43–45^, it remains unclear how this integration affects the reproducibility of the identified causal variants across independent studies. Ensuring that a putative causal variant is reproducible across different studies is crucial for its confident use in downstream causal inference and functional validation. With the availability of larger molecular QTL resources and the rapid advances in S2O models, we propose a generalized framework to utilize various deep genomic models to enhance genetic fine-mapping and systematically evaluate how these new tools can help identify causal variants that replicate across studies. Using eQTL resources from Genotype Tissue Expression (GTEx)^46^, Multi Ancestry analysis of Gene Expression (MAGE)^47^, and Trans-Omics Precision Medicine (TOPMed)^48^, we assess various strategies for incorporating S2O models to improve the reproducibility of genetic fine-mapping.

From this analysis, we provide practical recommendations and resources to facilitate functionally informed fine-mapping with these powerful new models.

## Results

### Fine-mapping replication failure rate (RFR) in molecular QTLs

Previous studies have reported that statistical fine-mapping can exhibit poor calibration, leading to replication failure rates (RFRs)—in these studies, defined as the proportion of high-confidence variants identified in smaller, downsampled subsamples of a cohort that fail to replicate in the full dataset—that exceed those predicted by the underlying statistical model^49^. We investigated whether these findings, originally observed in genome-wide association studies (GWAS) of 10 quantitative traits from UK Biobank^37^, extend to molecular traits such as expression quantitative trait loci (eQTLs). We hypothesized that the availability of thousands of cis-acting molecular traits would enable a more detailed examination of fine-mapping RFRs, with a simpler genetic architecture than that of the complex traits.

To assess the reproducibility of fine-mapped eQTLs, we employed a discovery-replication framework using two independent eQTL datasets. In our framework, we define RFR as the percent of credible sets in the discovery set that have no variant replicated in any credible set of the same gene from the replication set (Fig. 1a). The discovery set consisted of 6,958 fine-mapped credible sets from GTEx blood eQTLs (v10, n = 800)^46^, while the replication set comprised of 54,000 credible sets from TOPMed blood eQTLs (v1, n = 6,454) as the “replication” set^48^ (Fig. 1a). Although we expect >98% of credible sets in GTEx blood would contain a variant to achieve genome-wide significance (P < 5×10⁻⁸) in TOPMed under the simplistic assumption of the same effect sizes between the two datasets (see Methods), we observed much poorer reproducibility of credible sets between GTEx and TOPMed. Out of 2,126 high-confidence GTEx blood eQTL credible sets, each composed of a single SNP (i.e., credible set size = 1) with > 95% posterior inclusion probability (PIP), 697 (32.8%) did not replicate in TOPMed blood eQTLs, meaning that these putative causal variants were not found in any corresponding TOPMed credible sets for the same gene, despite their high PIPs (Supplementary Table 1a).

**Figure 1.**
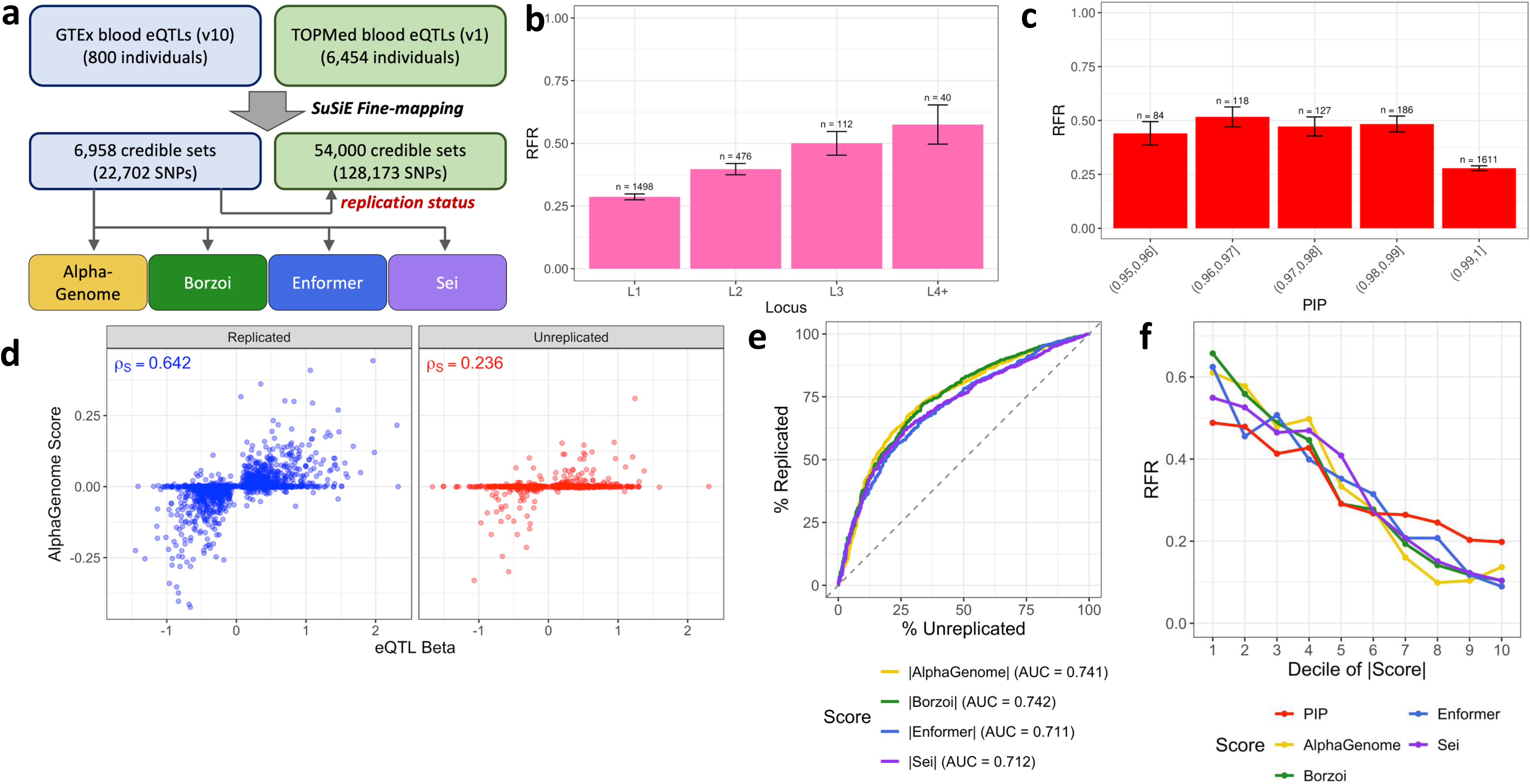
Reproducibility analysis of high-confidence GTEx blood eQTL credible sets. **(a)** Schematic diagram of replication failure rate (RFR) evaluation in blood eQTLs. SuSiE fine-mapped GTEx (v10) blood eQTL credible sets were used as “discovery” sets. The “replication status” of each variant within a credible set was determined by examining whether the variant is contained in the credible sets of a larger TOPMed (v1) blood eQTL analysis of the same gene. For each fine-mapped variant in GTEx blood eQTLs, their S2O scores were evaluated using AlphaGenome, Borzoi, Enformer, and Sei (targeting whole blood tissue) to examine their relationship with RFR. **(b)** Replication failure rate (RFR) of 2,126 high-confidence GTEx (v10) blood eQTL credible sets composed of a single SNP with PIP > 95%, stratified by the order of credible sets in SuSiE’s IBSS (Iterative Bayesian Stepwise Selection) algorithm. n indicates the number of credible sets in the category, and the error bars represent estimated standard errors. **(c)** Same as (b) but stratified by the PIP values. **(d)** Spearman correlation between eQTL effect sizes and AlphaGenome single track scores for 2,126 high-confidence (PIP > 95%) GTEx blood eQTLs, separated by their replication status in TOPMed blood eQTLs. **(e)** ROC curve of AlphaGenome, Borzoi, Enformer, and Sei single track scores in predicting the replication status of 2,126 high-confidence (PIP > 95%) GTEx blood eQTLs. **(f)** Replication failure rate (RFR) of 2,126 high-confidence GTEx (v10) blood eQTL credible sets (PIP > 95%), stratified by the deciles of PIP and single track S2O scores.

When we categorized these credible sets by their order in SuSiE’s forward selection algorithm within a locus, we observed RFRs of 28.6%, 39.7%, 50.0%, and 57.5% in primary (L1), secondary (L2), tertiary (L3), and higher-order (L4, L5, L6) credible sets, respectively (Fig. 1b, Supplementary Table 1b). This indicates that non-primary credible sets are less likely to replicate than primary credible sets, likely because of stronger confounding effect due to linkage equilibrium, weaker effect size from the putative causal variants, and/or inaccurate convergence in SuSiE’s IBSS (Iterative Bayesian Stepwise Selection) algorithm. Even when we applied a stricter PIP threshold (> 99%) to 1,611 GTEx credible sets, the RFRs only slightly decreased to 24.0%, 35.0%, 45.2%, and 50.0% respectively. These RFRs remain substantially higher than predicted by earlier simulation studies (Fig. 1c, Extended Data Fig. 1a, Supplementary Table 1c-d) and align with previous findings of higher-than-expected RFRs in fine-mapping studies^37^.

### Recent deep genomic models help improve fine-mapping reproducibility

While incorporating functional annotations into fine-mapping has been reported to enhance its power^43–45^, its impact on the reproducibility of fine-mapped loci remains unclear. We investigated how recent deep S2O models, such as AlphaGenome^24^, Borzoi^25^, Enformer^26^, and Sei^27^, improve fine-mapping reproducibility.

First, as described above, we focused on 2,126 putative causal SNPs in high-confidence (PIP > 95%) GTEx blood eQTLs. We then examined the association between their predicted single track S2O model scores (aimed at predicting their transcriptional changes in blood tissue, see Methods) and their eQTL effect sizes, as well as their fine-mapping reproducibility.

Consistent with previous findings^24^, we observed a strong Spearman’s correlation (ρ_S_ = 0.534) between AlphaGenome scores and eQTL effect sizes. Furthermore, when we examined TOPMed-replicated SNPs, we found a much stronger correlation (ρ_S_ = 0.642) compared to the TOPMed-unreplicated SNPs (ρ_S_ = 0.236) (Fig. 1d). Assuming that all TOPMed-replicated SNPs are causal and that AlphaGenome scores of noncausal variants are uncorrelated with eQTL effect sizes, we estimated that 23% of the 2,126 SNPs are noncausal (see Methods), which is much lower than the 32.7% observed for RFR. This suggests that a substantial fraction of RFR represents false negatives in the replication set (i.e., TOPMed) and RFR should be interpreted as a loose upper bound of false positives in fine-mapping. Borzoi, Enformer, and Sei scores showed similar trends but with weaker correlations than AlphaGenome (Extended Data Fig. 1b-d). The estimated proportions of non-causal variants were similar across Borzoi (21%), Enformer (24%), and Sei (23%).

TOPMed-replicated variants consistently exhibited stronger magnitudes of predicted S2O model scores than unreplicated variants across all four methods (Extended Data Fig. 1e), regardless of the directionality of these scores. When using only the magnitude of each S2O model score to classify the TOPMed replication status of each variant, the area under the ROC curve (auROC) ranged from 0.71 to 0.74 (Fig. 1e). We stratified the 2,126 high-confidence SNPs into 10 bins based on their S2O model scores and PIPs. Variants in the highest decile of S2O scores achieved a RFR of 9.0%-13.6% across the four methods, whereas variants with the highest PIP decile still showed a high RFR of 21.7% (Fig. 1f, Supplementary Table 1e). These findings suggest that S2O model scores offer additional information to purely statistical fine-mapping for improving the reproducibility of putative causal variants within fine-mapped loci. Moreover, when PIP and S2O scores are combined together, we observed that RFR can be further reduced. For example, when classifying each SNP to high/low S2O model or PIP scores based on their median values, we observed 13-15% RFR when both S2O model and PIP scores are low, and 56-60% RFR when both scores are high (Extended Data Fig. 1f).

### Sequence-to-omics models help prioritize replicable fine-mapped variants

We extended our fine-mapping reproducibility analysis beyond the high-confidence credible sets. For 6,958 GTEx blood eQTL credible sets with 10 or fewer SNPs, we assessed replication in TOPMed blood eQTL credible sets for the same gene. The replication failure rate (RFR) was determined as the proportion of GTEx credible sets with no variants replicated in TOPMed credible sets. RFRs remained high across all credible set sizes, ranging from 30.6% to 42.0% (Fig. 2a, Supplementary Table 2a). Primary (L1) credible sets had substantially lower RFRs (33.6%) compared to non-primary sets (47.0%), suggesting fine-mapping is less reproducible in complex loci with multiple putative causal variants (Fig. 2b, Supplementary Table 2b). Dividing credible sets into 10 bins according to the largest absolute S2O model score among their SNPs showed significantly lower RFRs in sets with high absolute S2O scores across all four models (Fig. 2c, Extended Data Fig. 2a-c, Supplementary Table 2c). For instance, credible sets in the highest decile of absolute S2O scores had 3.1-4.2 times lower RFRs than those in the lowest decile. In contrast, the highest decile of PIPs showed only 1.6 times lower RFR than the lowest decile of PIPs (Extended Data Fig. 2d, Supplementary Table 2c), indicating that S2O models are more informative than PIPs for prioritizing reproducible credible sets.

**Figure 2.**
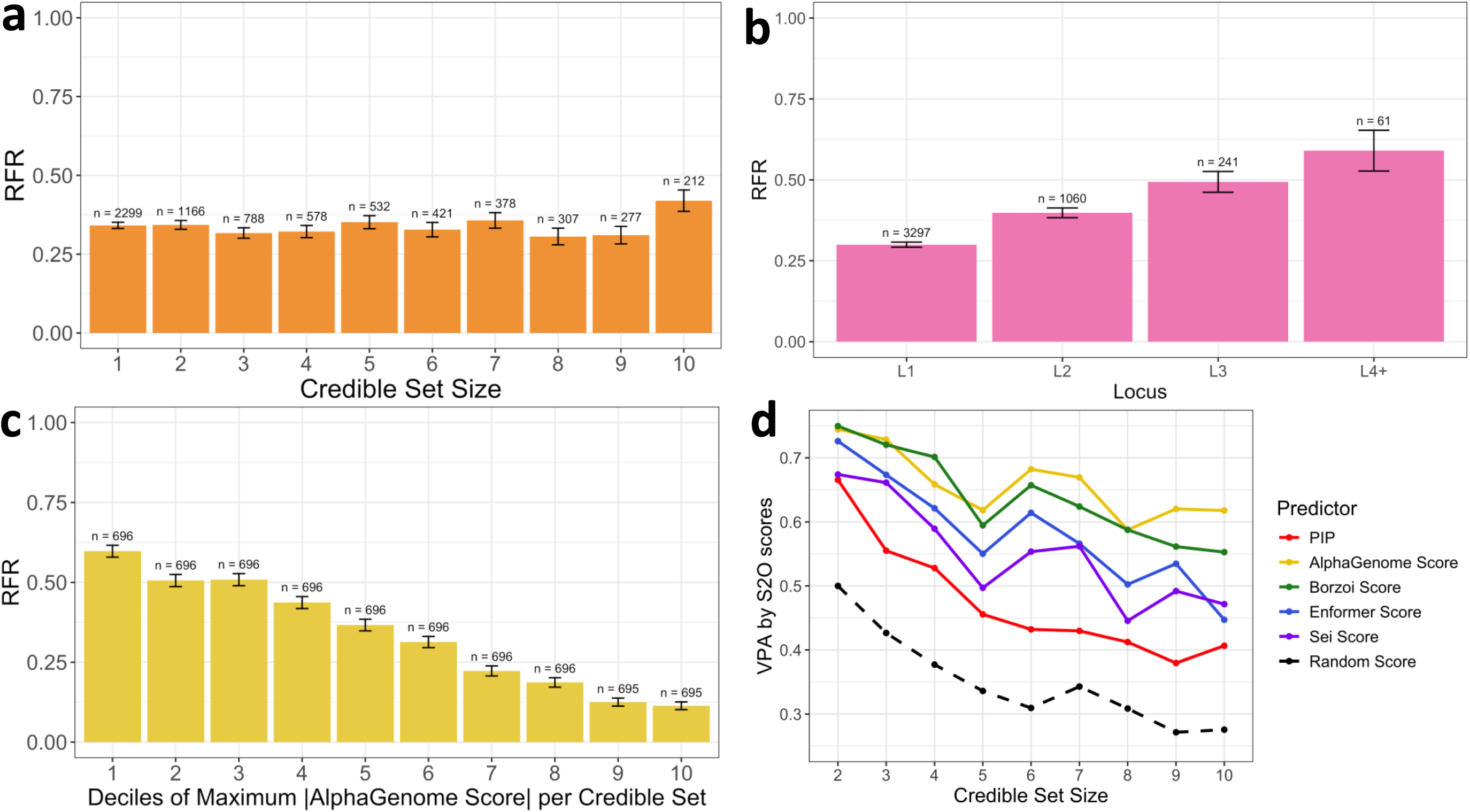
Reproducibility analysis of all GTEx blood eQTL credible sets with size ≤ 10. **(a)** Replication failure rate (RFR) of 6,958 GTEx (v10) blood eQTL credible sets with size 10 or less, stratified by SuSiE credible set size. n indicates the number of credible sets in the category, and the error bars represent estimated standard errors. **(b)** Same as (a) but stratified by the order of credible sets in SuSiE’s IBSS (Iterative Bayesian Stepwise Selection) algorithm. **(c)** Same as (a) but stratified by deciles of the maximum absolute AlphaGenome single track score per credible set. **(d)** Variant prioritization accuracy (VPA) of prioritizing replicated variants as the top variant, stratified by credible set size, comparing PIP and S2O single track scores. Random scores are used as a baseline.

Next, we evaluated the ability of each method to prioritize putative causal variants within a credible set, defining variant prioritization accuracy (VPA) as the fraction of credible sets for which a replicated variant was ranked as the top-scoring variant (see Methods). S2O model scores consistently showed higher VPA than PIPs across all credible set sizes (Fig. 2d, Supplementary Table 3a), with AlphaGenome and Borzoi performing best among all models. Similar trends were observed when stratifying credible sets by primary (L1), secondary (L2), tertiary (L3), and higher-order (L4+) loci (Extended Data Fig. 2e, Supplementary Table 3b). These findings indicate that S2O model scores not only reduce RFRs for each credible set but also enhance the reproducibility of the most likely causal variants within a given credible set. When credible sets were stratified by their maximum PIP, PIP-based VPA performed similarly to the best S2O models, AlphaGenome and Borzoi, for credible sets with a maximum PIP ≥ 0.8 (Extended Data Fig. 2f, Supplementary Table 3c). However, for credible sets with a maximum PIP ≤ 0.7, VPA was substantially poorer for PIP-based methods than for S2O models. This suggests that S2O models can offer complementary information to PIPs, thereby improving the reproducibility of fine-mapped variants when combining S2O models with statistical fine mapping.

### A generalized framework for functionally informed fine-mapping

We developed a general framework to systematically incorporate S2O models with fine-mapping to compute the functionally informed posterior inclusion probability (fiPIP) for each variant within a credible set. Our fiPIP framework shares similarity with previously developed methods such as PolyFun^43^ and expression modifier scores (EMS)^44^, but generalizes to an arbitrary set of quantitative scores from any functional prediction models (Fig. 3a). Our fiPIP framework, which includes S2O model score generation and fiPIP training and generation, is available as a public tool (see Code Availability).

**Figure 3.**
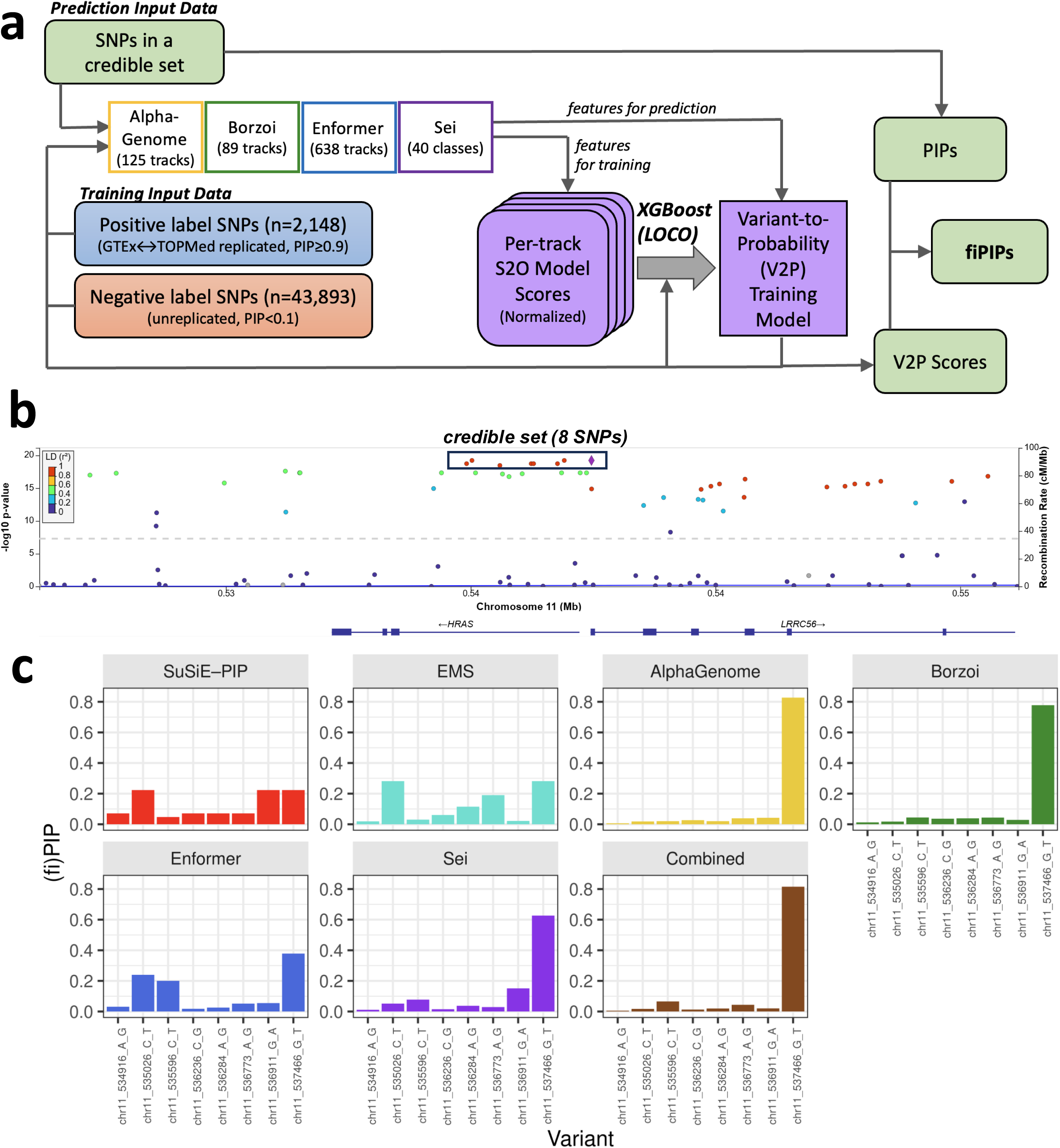
Overview and examples of functionally informed fine-mapping. **(a)** Schematic diagram of functionally informed fine-mapping framework. **(b)** LocusZoom plot of association signals for *LRRC56* cis-eQTLs, including 8 SNPs contained in the credible set. **(c)** Posterior inclusion probabilities (PIPs) from SuSiE fine-mapping, expression modifier scores (EMS), and functionally informed PIPs (fiPIPs) for *LRRC56* cis-eQTLs.

Our fiPIP framework first trains a variant-to-probability (V2P) model from a set of positive and negative SNPs, each annotated with a set of S2O model scores. Positive labels are chosen from confidently fine-mapped variants that show replication across multiple datasets with high PIP. Conversely, negative labels are selected from variants that do not replicate in other consortia and have a low PIP. In our experiments with GTEx and TOPMed blood eQTLs, we selected 2,148 positive and 43,898 negative label SNPs based on these criteria (see Methods). Using selected S2O model scores from these labels, V2P models are trained with XGBoost^50^ classifiers in a leave-one-chromosome-out (LOCO) manner to prevent overfitting (see Methods). For each credible set, fiPIPs can be computed by linearly reweighting original PIPs based on the V2P classifier scores (see Methods).

To illustrate fiPIP’s effectiveness, we examined a GTEx blood eQTL for *LRRC56*. The LocusZoom^51^ plot (Fig. 3b) for this locus shows a long block of linkage disequilibrium (LD) with over 50 genome-wide significant variants. While PIPs from SuSiE fine-mapping identified 8 SNPs within the 95% credible set, with PIPs ranging from 0.05 to 0.22, the fiPIPs pinpointed a single SNP (chr11_537466_G_T, rs112360405) with a much higher posterior probability (Fig. 3c, Supplementary Table 4). This SNP is located near the *LRRC56* promoter and was particularly highlighted when AlphaGenome and Borzoi were used or when all four S2O models were combined.

### Functionally informed fine-mapping prioritizes replicable variants with higher confidence

To systematically evaluate the benefits of fiPIP and compare the performance of different S2O models, we focused on 1,166 GTEx blood eQTL credible sets that contained exactly two SNPs. Among these, 595 credible sets had only one SNP replicated in the TOPMed blood eQTL credible sets, with the other SNP not being replicated. We then evaluated how effectively different functionally informed fine-mapping methods prioritized the replicated variant with a higher fiPIP.

We observed that the accuracy of PIPs from SuSiE statistical fine-mapping, without the use of S2O model scores, in prioritizing the replicated variant was 66.6%. In contrast, fiPIPs demonstrated significantly higher accuracy (73.8-79.3%). Among the four S2O models, the fiPIPs utilizing AlphaGenome (77.8%) and Borzoi (78.0%) achieved the highest individual accuracy, with the combined fiPIP reaching the highest overall accuracy of 79.3% (Fig. 4a, Extended Data Fig. 3a-3g, Supplementary Table 5). It is noteworthy that the previously published method EMS^44^, based on Basenji^52,53^ with a random forest classifier, also showed high accuracy (76.1%) similar to the Enformer-based fiPIP but lower than the AlphaGenome-and Borzoi-based fiPIPs. These findings suggest that the accuracy of individual S2O models is the primary determinant of fiPIP accuracy, rather than the specific architecture of the functionally informed fine-mapping framework.

**Figure 4.**
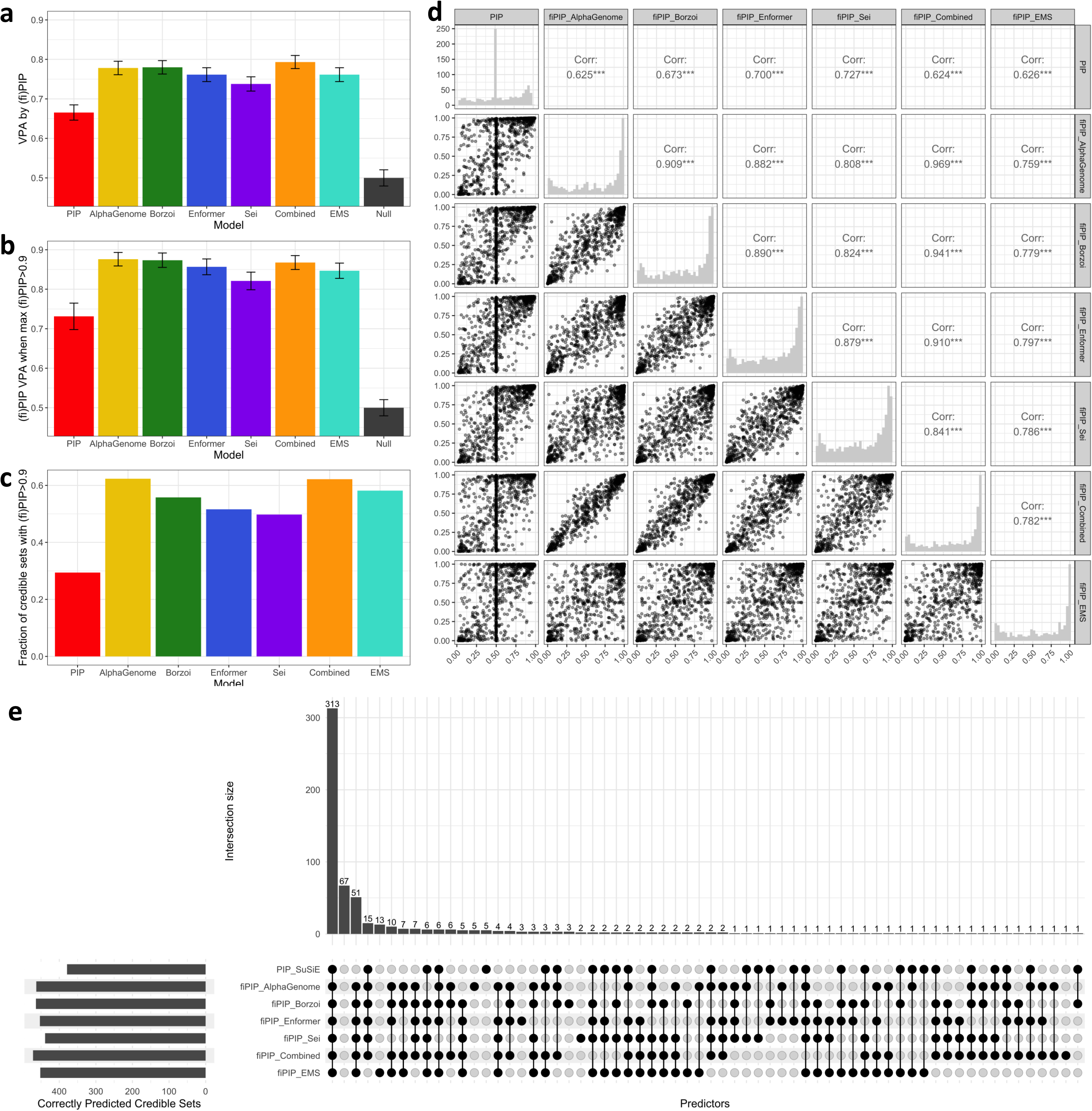
Evaluation of functionally informed fine-mapping for GTEx blood eQTL credible sets with size 2. **(a)** Variant prioritization accuracy (VPA) of SuSiE PIP and functionally informed PIPs (fiPIPs) across different S2O models for 595 credible sets with size 2 where only one SNP is replicated. **(b)** VPA of (fi)PIPs for size 2 credible sets containing a SNP with (fi)PIP ≥ 0.9. **(c)** Fraction of the 595 size 2 credible sets containing a SNP with (fi)PIP ≥ 0.9. **(d)** Pairwise scatterplots between (fi)PIPs of replicated SNPs across different methods for 595 credible sets of size 2. Diagonal plots show the histogram of the (fi)PIP values, and upper-diagonal panels summarize the pairwise Pearson’s correlation between methods. **(e)** Upset plot showing the distribution of credible sets that correctly prioritized the replicated variants across SuSiE PIP and different fiPIPs among the 595 credible sets of size 2.

More importantly, fiPIPs dramatically improve the power and accuracy of identifying confidently prioritized variants compared to conventional SuSiE-only fine-mapping. For instance, among 595 credible sets of size 2 with only one replicated SNP, only 175 (29.4%) were confidently prioritized by SuSiE-only fine-mapping (defined as having a highest posterior probability ≥ 0.9). Of these, only 128 (73.1%) correctly prioritized the TOPMed-replicated variant as the top variant. In contrast, the AlphaGenome fiPIP identified 371 (62.4%) credible sets as confidently prioritized, with 325 (87.6%) of them accurately prioritizing TOPMed-replicated variants (Fig. 4b-4c, Extended Data Fig. 3h-3n, Supplementary Table 5). The other three S2O models also demonstrated much greater power (49.7-55.8%) to confidently prioritize the TOPMed-replicated variant in the credible set with higher accuracy (82.1-87.3%) than SuSiE-only fine-mapping. Combining all four S2O models also achieved high power (62.1%) and accuracy (86.8%), similar to the best-performing AlphaGenome method. Pairwise comparisons of replicated variants between methods show that SuSiE-only fine-mapping often cannot distinguish between the two SNPs, likely due to strong pairwise LD, while fiPIPs distinguish most of them by leveraging S2O model scores (Fig. 4d). It is also noteworthy that in 380 (64%) credible sets, fiPIPs from all tested S2O models and PIPs from SuSiE fine-mapping consistently prioritized the same replicated SNP. However, in 67 (18%) of them, the unreplicated SNP was prioritized over the replicated SNP (Fig. 4e), which can be partially explained by inaccurate replication-based ground truth labels as mentioned earlier.

### Detailed evaluation across various configurations of functionally informed fine-mapping

Next, we extended our evaluation of functionally informed fine-mapping to all credible sets, focusing on VPA. Across the four S2O models, we observed that across credible set sizes, within-locus orders, and maximum PIP values, fiPIPs improved VPA compared to using either S2O models or PIPs alone (Fig. 5a, Extended Data Fig. 4a-4b, Supplementary Table 6a-c). These improvements were twofold: first, when a single blood track (see Methods) was used, VPA improved when S2O models and PIPs were combined into fiPIPs compared to using the S2O model alone by 1.4-4.9% (Fig. 5b, Supplementary Table 6d).

**Figure 5.**
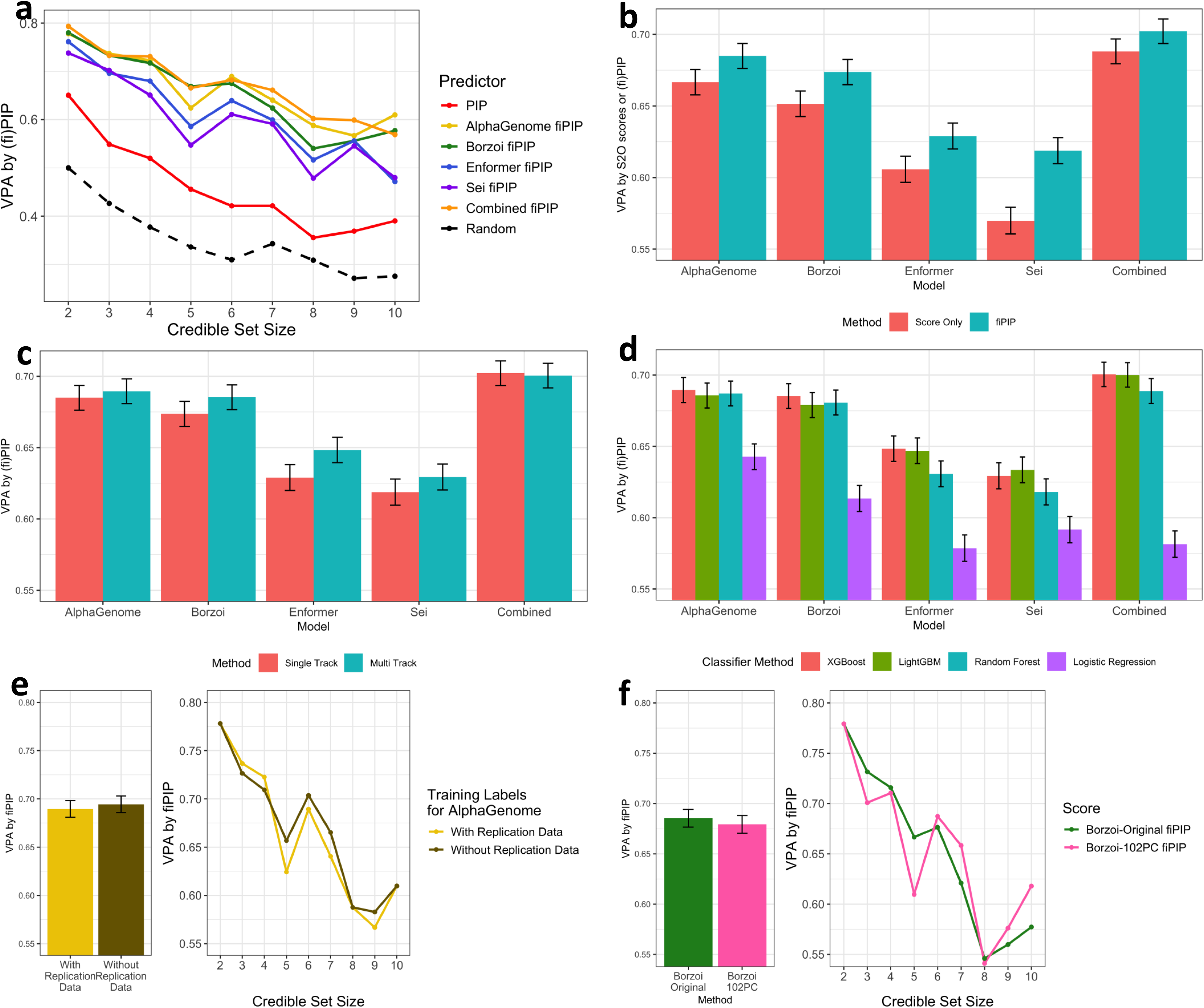
Detailed evaluation of functionally informed fine-mapping using GTEx blood eQTLs. **(a)** Variant prioritization accuracy (VPA) of prioritizing replicated variants as the top variant using functionally informed posterior inclusion probabilities (fiPIPs) trained by multitrack S2O models, stratified by credible set size. Five fiPIPs based on AlphaGenome, Borzoi, Enformer, Sei, and a combined model using scores from all four S2O models were compared, in addition to PIP and random scores. **(b)** VPA comparison between variant-to-probability (V2P) scores trained from S2O models and fiPIPs that combine V2P scores with PIP, using single track S2O models. AlphaGenome, Borzoi, Enformer, Sei, and combined models were evaluated. **(c)** VPA comparison between single track and multitrack fiPIPs across five S2O models – AlphaGenome, Borzoi, Enformer, Sei, and combined models. **(d)** VPA comparing multitrack fiPIPs based on V2P scores trained by different classifiers: XGBoost, LightGBM, random forest, and logistic regression. AlphaGenome, Borzoi, Enformer, Sei, and combined models were evaluated. **(e)** VPA comparing multitrack AlphaGenome fiPIPs using different sets of training labels. “With Replication Data” represents the original labels used in this paper, taking positive labels from replicated variants with high PIP, and negative labels from unreplicated variants with low PIP (see Methods). “Without Replication Data” uses alternative training labels that do not require a replication dataset, by taking positive labels from high PIP variants, and negative labels from variants not included in GTEx credible sets of any tissue. **(f)** VPA comparing between multitrack Borzoi fiPIPs used in this paper and fiPIPs trained by the 102-dimensional Borzoi annotation set (Borzoi-102PC) designed for the Sniff method.

Second, utilizing multiple (40-638) track scores further improved VPA compared to using a single blood track alone, but the improvement was much more modest, ranging from 0.2%-1.0% (Fig. 5c, Supplementary Table 6e) across all models. In addition, using all 892 available tracks from multiple S2O models in a combined model showed higher VPA compared to the two best-performing models on their own, AlphaGenome and Borzoi (Fig. 5a-d, Supplementary Table 6a, d-f).

We also evaluated whether the choice of classifiers to train V2P models substantially affects the performance of functionally informed fine-mapping. When XGBoost^50^ was replaced with LightGBM^54^ for training, the VPAs were very similar (Fig. 5d, Supplementary Table 6f). However, training models with logistic regression engendered considerably poorer performing VPAs than either gradient boosting method. Training models with random forest engendered VPAs that were on average higher than logistic regression but lower than the gradient-boosting methods.

Given the challenge of consistently obtaining replication status across multiple eQTL datasets for positive and negative labels, we assessed how training models without TOPMed replication dataset impacts the accuracy of fiPIPs. Following prior studies^44^, we chose positive labels from credible sets that contained only a single SNP with a high PIP (≥ 0.9), without considering their replication status. Negative labels were chosen from variants within a 50kb distance of transcription start sites (TSS) that were not present in any credible sets across 49 GTEx tissues. This alternative method for selecting training labels did not substantially lower the fiPIP VPAs for AlphaGenome (Fig. 5e, Supplementary Table 7a-b), suggesting that having replication status for positive and negative labels may not be crucial for constructing effective V2P training models for accurate functionally informed fine-mapping.

Finally, we evaluated whether the recently developed Borzoi-informed fine-mapping resource^45^, designed for Sniff, could be incorporated into our functionally informed fine-mapping framework. Although Sniff is a modification of the PolyFun fine-mapping method targeting complex phenotypes, we can leverage its underlying 102 principal component (PC)-dimensional Borzoi annotation set (Borzoi-102PC) within our fiPIP framework. This is achieved by replacing our multitrack function annotation set with Borzoi-102PC, which comprises principal components across 7,611 functional genomic coverage tracks, including RNA-, ChIP-, CAGE-, DNAse-, and ATAC-seq. Despite Borzoi-102PC not being gene-specific, we observed that fiPIPs based on Borzoi-102PC showed comparable variant prioritization accuracy (VPA) to our original Borzoi-based fiPIPs (Fig. 5f, Supplementary Table 8a-b). Given that Borzoi-102PC annotations are already precomputed for 19.53 million common and low-frequency variants, functionally informed fine-mapping using Borzoi-102PC scores offers a convenient alternative to our original approach, which requires computing S2O model scores for each variant.

### Analysis of lymphoblastoid cell line eQTLs

We also assessed RFRs and VPAs in additional eQTL datasets from lymphoblastoid cell lines (LCL) using GTEx (v10) and MAGE, anticipating lower replication due to smaller sample sizes (n = 326 for GTEx and n = 731 for MAGE). Focusing on high-confidence credible sets (PIP ≥ 0.95, credible set size 1), we observed an RFR of 46.8% in LCL eQTLs, which is indeed higher than blood eQTLs (32.8%). Nevertheless, similar to blood eQTLs, we observed highest Spearman correlation between AlphaGenome scores and eQTL effect sizes (ρ_S_ = 0.642) in the replicated high-confidence credible set (Fig. 6a), followed by Borzoi (ρ_S_ = 0.558), Enformer (ρ_S_ = 0.378), and Sei (ρ_S_ = 0.264) (Fig. 6a, Extended Data Fig. 5a-c). The estimated proportions of non-causal variants in the high-confidence credible sets ranged from 27%-33% across the S2O models (see Methods). Across all credible sets with size 10 or less, for single track scores without PIP integration, AlphaGenome and Borzoi had the highest VPAs at 0.715 and 0.701 respectively, followed by Enformer (0.671) and Sei (0.627), which were all significantly higher than PIP alone (0.561) (Fig. 6b, Supplementary Table 9a). When PIPs and multitrack scores were combined into fiPIPs, VPAs increased across all methods: AlphaGenome (0.725), Borzoi (0.723), Enformer (0.691), and Sei (0.642) (Fig. 6c, Supplementary Table 9b). These findings are consistent with our blood eQTL observations, suggesting that AlphaGenome and Borzoi are consistently better for enhancing the reproducibility of functionally informed fine-mapping across different tissues and datasets.

**Figure 6.**
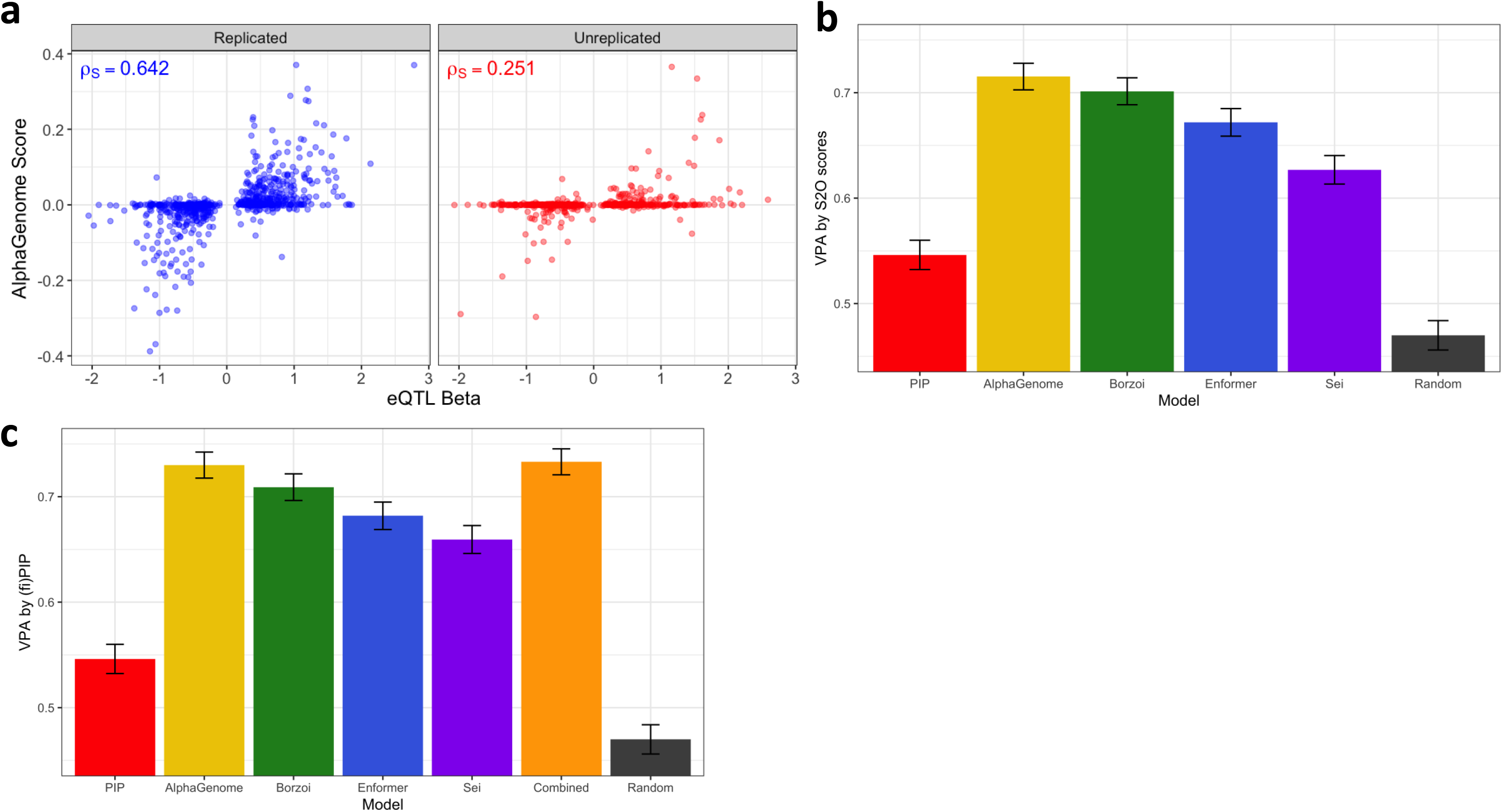
Evaluation using GTEx LCL eQTLs. **(a)** Spearman correlation between eQTL effect sizes and AlphaGenome single track scores for 1,364 high-confidence (PIP > 95%) GTEx LCL eQTLs, separated by their replication status in MAGE LCL eQTLs. **(b)** Variant prioritization accuracy (VPA) of single track S2O model scores for 6,673 GTEx LCL credible sets of size 2 to 10, using replication status in MAGE LCL eQTLs. **(c)** Variant prioritization accuracy (VPA) of multitrack fiPIPs for 6,673 GTEx LCL credible sets of size 2 to 10, using replication status in MAGE LCL eQTLs.

### Practical considerations in utilizing sequence-to-omics models for fine-mapping

During our experiment evaluating various S2O models for fine-mapping, we identified specific technical characteristics that can guide users in selecting a model for functionally informed fine-mapping. For instance, the publicly available Enformer resource^26^ uses an unconventional standard for identifying variants, which can lead to technical artifacts if not carefully processed. We discovered this issue by comparing the eQTL effect sizes of high-confidence GTEx blood eQTL credible sets. Without correcting for potential allele flipping, we observed much stronger directional inconsistencies between Enformer scores and eQTL effect sizes (Extended Data Fig. 6a). The vast majority of these flipped alleles have GTEx allele frequencies > 0.5, suggesting that minor-major allele encoding was inadvertently used instead of reference-alternate allele encoding in the published resource (Extended Data Fig. 6b). While our results are based on corrected allelic representations, other published studies also report and/or account for similar patterns of directional mixtures^33,34,55–57^, suggesting the importance of considering this technical aspect when using precomputed Enformer scores for common variants.

When performing functionally informed fine-mapping at scale, it is important to consider the heterogeneous computational costs associated with generating S2O model scores. We observed that without GPU resources, computing Borzoi scores is time-consuming (244.5 seconds/variant), whereas Sei is significantly more computationally scalable (4.6 seconds/variant). The computational cost for generating AlphaGenome scores is relatively low (4.3 seconds/variant), but these are currently only available via an API. If precomputed S2O model scores are available across the vast majority of common and low-frequency variants, such as in Enformer or Borzoi-102PC, it will greatly facilitate the utility of the corresponding S2O model for functionally informed fine-mapping.

## Conclusion

In this study, we investigated the utility of existing sequence-to-omics (S2O) models in enhancing the reproducibility of fine-mapping by leveraging multiple large-scale eQTL resources. Our comprehensive evaluation across datasets and methodologies consistently demonstrated that functionally informed fine-mapping, when augmented by S2O models, substantially improves variant prioritization accuracies (VPAs) of reproducible variants. Notably, more recent methods such as AlphaGenome and Borzoi substantially outperformed earlier models, highlighting the rapid advancements in this field. These findings underscore the potential of integrating S2O models to refine the identification of causal variants in genetic studies.

Furthermore, our observations revealed that functionally informed fine-mapping leads to the production of more credible sets containing highly confident (PIP > 0.9) putative causal variants. Linkage disequilibrium (LD) often hinders confident pinpointing of causal variants based solely on statistical evidence. Our results show that S2O models provide additional information to accurately prioritize putative causal variants that are subsequently and independently reproduced in other studies.

Our study primarily focused only on SNPs because some S2O models are prone to systematic bias in predicting the impact of insertions, deletions, and other complex variants^25^. However, given the observations that indels and tandem repeats explain a large proportion heritability of molecular traits^58^, it is important to comprehensively incorporate a wider range of variants to comprehensively evaluate the benefit of S2O models in fine-mapping in the future, especially given that newer methods are reported to more accurately model the impact of indels and complex variants^59^.

Our study also focused on evaluating the replication failure rates (RFRs) or variant prioritization accuracy (VPAs) of existing credible sets rather than generating better credible sets by incorporating S2O models as priors for functionally informed fine-mapping. While the assignment of different priors across all SNPs during fine-mapping, as implemented in methods like PolyFun^43^ and Sniff^45^, offers a path to more accurate credible sets in principle, its practical application is contingent on the availability of S2O model scores for nearly all variants. Therefore, a broader accessibility of precomputed S2O model scores is essential to facilitate the adoption of such integrated, functionally informed fine-mapping frameworks. More significantly, if precomputed S2O model scores are available for all 9 billion possible SNPs, it could dramatically enhance our capacity to include rare functional variants in genetic fine-mapping. As S2O models continue to evolve, their utility in fine-mapping is expected to grow. To maximize their practical impact, it is crucial to make necessary resources available to the community, reduce computational burdens, and ensure software tools are easily adaptable to incorporate newer S2O models.

## Methods

### eQTL resources

The eQTL credible set and fine-mapping resources used in this experiment come from publicly released credible sets from whole blood eQTLs from GTEx v10^46^ (n = 800 individuals, 6,958 credible sets, 22,702 SNPs) and TOPMed^48^ (n = 6,454 individuals, 54,000 credible sets, 128,173 SNPs). We keep only SNPs from the eQTL credible sets; all indels were removed, and PIPs within a credible set were readjusted to sum to 1 when compared with other fiPIPs. For our experiments, we only consider credible sets with a maximum size of ten. For our Epstein-Barr virus (EBV)-transformed lymphoblastoid (LCL) eQTL experiment, LCL eQTLs from GTEx v10^46^ (n = 326 individuals, 4,862 credible sets, 17,906 SNPs) and MAGE^47^ (n = 731 individuals, 9,237 credible sets, 30,636 SNPs) were used.

### Replication, replication failure rate (RFR), and variant prioritization accuracy (VPA)

We consider eQTL fine-mapping results across a set of genes 𝐺. For each gene 𝑔 ∈ 𝐺, let 𝐶*_g_* denote the collection of credible sets and |*𝐶_g_*| the number of credible sets for that gene. For a credible set 𝑐*_g_* ∈ 𝐶*_g_*, we define 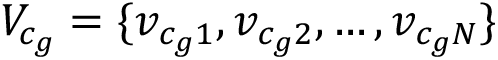 to be the set of SNPs in the credible set where 𝑁 = |𝑉*_cg_* |. Given a discovery set 𝐷 and replication set 𝑅, we determine a SNP in the discovery consortium, 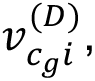 to be replicated if there exists a SNP,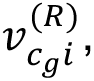 in the replication consortium such that both credible sets 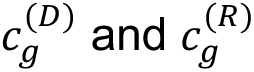 map to the same gene via Ensembl ID and belong to the same tissue. The variant replication status 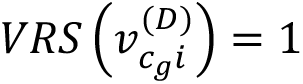 for replicated variants and 0 otherwise. We determine the variant replication status of SNPs in the replication consortium, 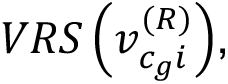 in a reciprocal manner.

A credible set 𝑐*_g_* is considered replicated if 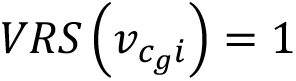 for at least one of its 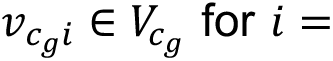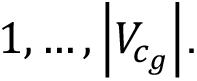 We define credible set replication status 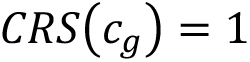 for replicated credible sets and 0 otherwise.

We define replication failure rate (RFR) on the credible set level as 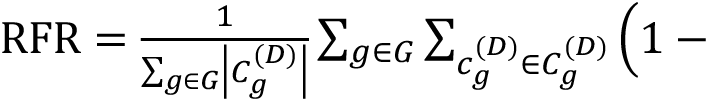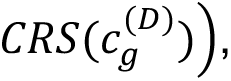 the proportion of unreplicated credible sets in the discovery set. To quantify how well a S2O model prioritizes replicated SNPs, we define variant prioritization accuracy (VPA) as the proportion of replicated variants among the best-scoring variants within each credible set for a given score metric. More specifically, for each credible set 𝑐*_g_*, we identify 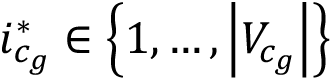 as the index of the variant with maximum score, e.g., PIP or S2O model scores, and we then define VPA as 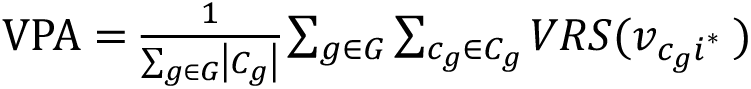 for the score metric, after excluding credible sets where 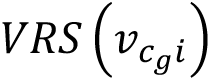 is identical, i.e. all 0 or all 1, across all variants within the credible set.

In our evaluation for whole blood eQTLs, we consider GTEx eQTLs as our discovery set, and TOPMed eQTLs as our replication set. For LCL eQTLs, we consider GTEx eQTLs as our discovery set, and MAGE eQTLs as our replication set. All indels and complex variants were removed when constructing 𝑉_cg_, and credible sets with |𝑉_cg_ | > 10 were also removed from the evaluation.

The simplistic estimate of statistical power to reach genome-wide significance in the replication dataset was calculated by taking the marginal association statistics for each variant within the GTEx discovery credible set, adjusting the sample size assuming that the effect sizes remain the same, and counting the proportion of credible sets where p-value of one of its SNPs reach the genome-wide significance.

The proportion of non-causal variants in high-confidence (PIP ≥ 0.95, credible set size 1) credible sets was estimated in the following way. The Spearman correlation between each S2O model scores and eQTL effect sizes among replicated variants and all variants are computed as 𝜌*_s,replicated_*:and 𝜌*_S,all_* respectively. Assuming that replicated variants are all causal variants and truly non-causal variants have zero correlation between S2O model scores and eQTL effect sizes, the proportion of non-causal variants among the high-confidence credible sets is approximated as 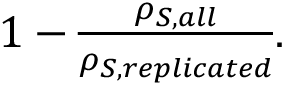 This simplistic approximation resulted in similar estimates of non-causal variants across the 4 different S2O models.

### SNP effect predictions for sequence-to-omics models

We use quantitative scores from three Transformer-CNN hybrid models AlphaGenome^24^, Borzoi^25^, and Enformer^26^ and one CNN model Sei^27^ to evaluate the role of S2O models in identifying casual SNPs. These models take as input a reference and alternative allele DNA sequence centered on the SNP of interest and generate predictions for molecular and epigenomic readouts (tracks). Our study investigates the utility of both single track and multitrack S2O scores. Single track S2O scores use one or a small number of cherry-picked available tracks from an S2O model: either only a single selected track or an average over a small number of tracks to ultimately generate one track. When possible, we cherry-picked tracks associated with gene expression of the tissue of interest (in our case, either whole blood or LCL depending on the experiment). For multitrack S2O scores, we instead used as many tracks as the S2O model provided that were associated with gene expression, prioritizing RNA-seq when possible. Each of these tracks were then used as predictors in the multitrack V2P generation process. In the single track V2P generation process, we use the single track S2O score as one predictor. For each method, we prioritize generating and using scores in a way close to the suggestions of the authors or using pre-generated scores when available.

### AlphaGenome scores generation

AlphaGenome scores for each SNP were generated using the AlphaGenome^24^ API. AlphaGenome is a hybrid Transformer-CNN model that takes up to 1 Mb of DNA sequence as input, predicts molecular regulatory features, and quantifies the effects of SNPs by comparing mutated vs. unmutated sequences. AlphaGenome output is of dimensions 𝑨 ∈ ℝ*^gA^*^×667^ where 𝑔_A_ is the total number of genes in the 1 Mb interval and 667 represents the number of total predictable RNA-seq tracks. Our AlphaGenome multitrack scores contain 125 tracks. There are 667 RNA-seq tracks that AlphaGenome can make predictions for; however, for both blood and LCL eQTLs, only 125 of the 667 tracks had no missing values generated for all SNPs in the discovery set. These 125 tracks include all the 54 GTEx RNA-seq tracks. The other 71 included tracks include assorted polyA plus RNA-seq tracks belonging to uber-anatomy, experimental factor, cell, and cell line ontologies. An explicit list of which tracks are used is available in Supplementary Table 10. The 524 other tracks each had at least 50% missing values for all 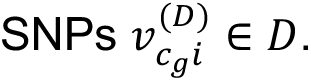 For each SNP 𝑣*_cg_i* belonging to credible set 𝑐_g_, we define its AlphaGenome score as 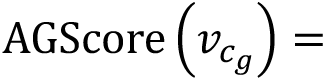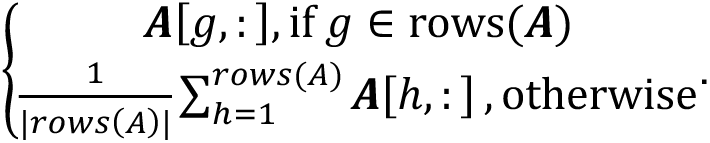 For blood eQTLs, the “UBERON:0013756 gtex Whole_Blood polyA plus RNA-seq” track was used for single track S2O scores. For LCL eQTLs, the “EFO:0000572 gtex Cells_EBV-transformed_lymphocytes polyA plus RNA-seq” track was used for single track S2O scores.

### Borzoi scores generation

The eQTL example notebook provided in the Borzoi^25^ repository was modified and used to generate Borzoi scores for each SNP in the discovery and replication consortia. Borzoi is a hybrid Transformer-CNN model that takes a 524 kb DNA sequence and predicts molecular regulatory features at 32 bp resolution, producing vectors of 524 kb / 32 bp = 16,384 bins for each track. In this study, we focus on the 89 GTEx RNA-seq tracks, predicting RNA-seq output for the 16,384 bins for each the reference and alternative allele, for each of 4 folds, leading to a separate reference allele tensor 𝑩_𝑹_ ∈ ℝ^4×16,384×89^ and alternative allele tensor 𝑩_𝑨_ ∈ ℝ^4×16,384×89^. To stabilize the variance and downweigh outliers, we apply the element-wise transformation 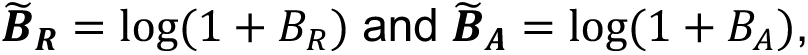 and then subsequently take the mean across the 4 folds to generate reduced 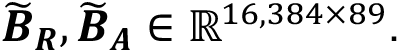 Then, we prioritize the 32 bp bins that have ≥ 50% overlap with exon(s) in a GENCODE release 41^60^ basic annotation GTF file by generating 𝑥 ∈ ℝ^16,384^, a {0, 1} vector with 0 values for bins not overlapping ≥ 50% with exons and 1 otherwise to generate 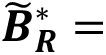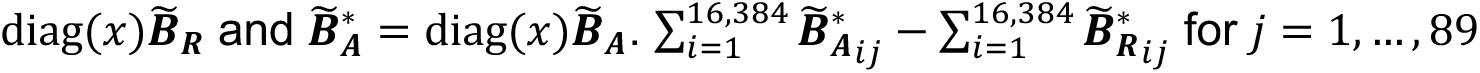 then produces a single-dimensional score BorzoiScore (𝑣*_cg_*) ∈ ℝ^89^ for each of the 89 GTEx RNA-seq tracks. We use the 89 GTEx RNA-seq tracks individually in the multitrack V2P score generation. To generate a single S2O score per SNP, we take the mean of the 2 GTEx blood + 1 GTEx LCL tracks for the blood eQTLs and just use the 1 GTEx LCL track score for the LCL eQTLs.

### Enformer scores generation

Enformer^26^ scores for each SNP were collected from pre-computed SNP effect predictions released by DeepMind. Enformer is a hybrid Transformer-CNN model that takes a 200 kb DNA sequence as input and predicts molecular regulatory features at 128 bp resolution. Pre-computed Enformer scores are of the dimensions 𝑬 ∈ ℝ^27,805,938×5,313^ as they have precomputed 5,313 tracks for 27,805,938 variants. 638 of these 5,313 tracks are CAGE tracks which are used for our multitrack V2P generation, eliciting EnformerScore (𝑣*_cg_*) ∈ ℝ^638^. Any credible set with at least one variant which was not present in the pre-computed Enformer score predictions was removed from downstream analyses. For the single track S2O scores, a mean of 5 CAGE tracks associated with blood + 1 CAGE track associated with LCLs was used for the blood eQTLs; and for the LCL eQTLs, the 1 CAGE track associated with LCLs was used.

### Sei scores generation

Sei^27^ scores were generated for each SNP using the variant effect prediction and sequence class prediction functions. Sei is a deep learning CNN framework that classifies genomic sequences into 40 sequence classes with each sequence class integrating predictions for 21,907 epigenomic profiles (transcription factor, histone marks, chromatin accessibility profiles, etc.). Each sequence class is treated as a track. Sei output for its epigenomic profile prediction for 𝑉 variants is of the dimensions 𝑺_1_ ∈ ℝ^v×21,907^. Epigenomic profile predictions 𝑺_1_ are then used as input for sequence class predictions which is of dimensions 𝑺_2_ ∈ ℝ^v×40^ as there are 40 sequence classes. Each sequence class is annotated with a functional pattern of regulatory activity across the genome. For the single track S2O scores, a single sequence class, HET6 Centromere, was used that had the maximum Spearman correlation (ρ_S_ = 0.340) between sequence class scores and eQTL effect sizes for likely causal variants (PIP ≥ 95%, credible set size 1) for the blood eQTLs (Supplementary Table 11). For multitrack V2P generation, all 40 sequence classes and their scores were used downstream, so we have SeiScore (𝑣_cg_) ∈ ℝ^40^.

### Variant-to-Probability (V2P) score generation and functionally informed PIP (fiPIP) framework

Our study uses scores from S2O models to assess their ability to prioritize putative causal variants in credible sets compared to results from statistical fine-mapping. First, we generate scores from S2O models as defined in the previous sections for all SNPs in all eQTL credible sets for both the discovery and replication sets. We then standardize the S2O scores for each track to have mean = 0 and standard deviation = 1 across all 𝑛 SNPs in the discovery and replication set together, e.g., a track 𝑤 ∈ ℝ*^n^* is transformed to 𝑧 by 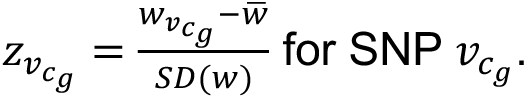 Next, to avoid overfitting in the evaluation, we use a leave-one-chromosome-out (LOCO) XGBoost^50^ classifier to generate sequence-to-omics variant-to-probability (V2P) scores for each SNP, denoted 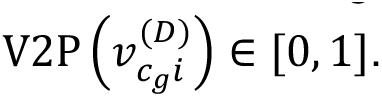 When making predictions for SNPs on one autosome, the classifier is trained only on SNPs from all other autosomes. We consider a SNP with greater V2P as more likely to be replicated. The positive labels for our XGBoost classifier are replicated SNPs from the GTEx consortium with PIP ≥ 0.9 and belonging to a singleton credible set of size 1, e.g., any replicated 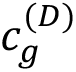 with only one SNP. The negative labels for our classifier are unreplicated SNPs with PIP ≤ 0.1. The predictors are the absolute values of the rescaled S2O scores, e.g. |Score(𝑣_cg_)|, where 𝑣_cg_ is an arbitrary SNP, and Score is one of AGScore, BorzoiScore, EnformerScore, or SeiScore, reflecting the assumption that the magnitude of regulatory impact is more relevant than its direction for replication prioritization. We test each S2O model separately using tracks defined in the previous section as well as a combined classifier that uses tracks from all of AlphaGenome, Borzoi, Enformer, and Sei together. Following PSS generation, we calculate fiPIPs for each SNP 𝑖 = 1,…, 𝑉 in a credible set 𝑐*_g_* by defining 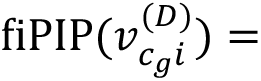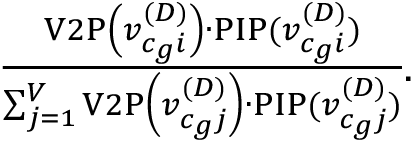

### Comparisons among XGBoost, LightGBM, random forest, and logistic regression

We use XGBoost^50^ as our classifier for our fiPIP framework, but we tested LightGBM^54^, random forest, and logistic regression as options for classifiers as well. Results comparing the four classifiers are in Fig. 5d. All classifier training, testing, and calculations were done in R. The xgboost^62^ R package version 1.7.11.1 was used. We trained gradient boosting models using the XGBoost framework with a binary logistic objective and optimized performance using the logarithmic loss metric. Each model was trained for 100 boosting rounds with a maximum tree depth of 3 and a learning rate of 0.1. Default xgboost settings were otherwise used. Random seeds were fixed for reproducibility.

Gradient boosting models were trained using the LightGBM framework via the lightgbm^63^ R package version 4.6.0 with a binary classification objective and evaluated using the binary log-loss metric. Models were trained for 100 boosting rounds with a learning rate of 0.1 and a maximum of 7 leaves per tree, which corresponds to a maximum tree depth of approximately 3. Default LightGBM settings were otherwise used.

Random forest classifiers were fit using the ranger^64^ package version 0.17.0 with the negative and positive label outcome encoded as a factor with levels 0 and 1 respectively, and predictors given by the absolute values of S2O scores. Models were trained with 300 trees, and the number of variables randomly sampled at each split was set to the square root of the total number of predictors. Variable importance was not computed, and the models were configured to output class probabilities. Default random forest settings from the ranger package were sued otherwise.

Logistic regression models were fit using the generalized linear model (GLM) framework with a binomial error distribution and a logit link function in the R stats package. Model coefficients were standardly estimated via maximum likelihood estimation, and predictions were obtained by applying the expit transformation to the fitted linear predictors.

### Summary of fiPIP experiments

Our experiments evaluate how sequence-to-omics (S2O) model scores can improve reproducibility in statistical genetic fine-mapping. We began by estimating baseline replication failure for high-confidence singleton credible sets (PIP ≥ 0.95) and, within these sets, tested whether single-track S2O scores correlate with eQTL effect sizes and distinguish replicated from unreplicated variants in TOPMed. We then broadened to all credible sets with ≤ 10 SNPs, re-estimating replication failure rates (RFRs) and assessing within-set prioritization via variant prioritization accuracy (VPA) for sets of size ≥ 2.

We next introduced the fiPIP framework to integrate functional predictions with SuSiE-only PIPs. We trained variant-to-probability (V2P) models on S2O annotations using replicated high-PIP singletons as positives and low-PIP unreplicated variants as negatives, applying a leave-one-chromosome-out (LOCO) model. fiPIPs were computed by reweighting SuSiE PIPs by V2P scores within each credible set and renormalizing, with both per-model implementations and a multi-model implementation.

To cleanly test whether adding functional information resolves ambiguous LD scenarios, we analyzed 1,166 blood credible sets containing exactly two SNPs, focusing on the 595 sets where only one SNP replicated. We compared SuSiE PIP and fiPIPs in terms of accuracy of identifying the replicated SNP and power to confidently prioritize that SNP (top (fi)PIP ≥ 0.9).

We performed a suite of sensitivity analyses. First, we evaluated whether using multiple tracks of gene expression (when available) from all available tissues improves VPA beyond a cherry-picked single track. Our choice of classifier was also tested by comparing XGBoost, LightGBM, random forest, and logistic regression. We then also evaluated if training could be successful without explicit replication labels, jettisoning the TOPMed-replication status from positive labels and redefining negatives as proximity-to-TSS variants absent from GTEx credible sets of all tissues. Finally, we swapped our multitrack Borzoi scores for the Sniff Borzoi-102PC resource to evaluate how those pre-computed scores fared in our fiPIP framework.

Lastly, to test transfer across tissues and smaller cohorts, we repeated core analyses in LCL eQTLs (GTEx as the discovery set, MAGE as the replication set). We measured baseline RFR for high-confidence singletons, correlations between S2O scores and effect sizes by replication status, and VPA for PIP, S2O scores, and fiPIPs across credible sets of size 2-10.

### Allele mismatch adjustment for Enformer scores

The SNPs in the credible sets were first lifted over from hg38 to hg19 using the LiftOver^61^ function of the UCSC Genome Web Browser. Then, SNPs from the credible sets were matched to the pre-computed Enformer scores. If a SNP from the credible set did not exactly match a SNP in the pre-computed scores, such as in the event of a reference and alternative allele swap, scores were multiplied by-1; however, sign has no effect on V2P generation. If there was not a direct match or a simple reference/alternative allele swap, the mean Enformer score was calculated across all variants with a matching chromosome and position.

## Data Availability

All GTEx eQTL summary statistics and fine-mapping results used in this paper can be downloaded publicly at GTEx Portal. The TOPMed eQTL summary statistics and fine-mapping results will be accessible in the TOPMed Genomic Summary results repository (dbGap phs001974). The MAGE eQTL summary statistics and fine-mapping results are publicly available in the corresponding Zenodo repository (https://zenodo.org/records/10535719). The input datasets to generate sequence-to-omics (S2O) model scores are accessible from the publications^24–27^. The expression modifier scores (EMS) across were requested to and obtained from the authors. All the relevant credible sets, S2O model scores, trained models, and fiPIP scores to reproduce our results will be publicly available through a Zenodo repository.

## Code Availability

The source code that implements our functionally informed fine-mapping pipeline is available at https://github.com/statgen/fipip. Source code for analysis and figures to reproduce our results will be available at https://github.com/statgen/fipip_manuscript. XGBoost variant-to-probability (V2P) models were trained in R using the xgboost^62^ R package version 1.7.11.1. For our classifier comparison, LightGBM V2P models were trained using the lightgbm^63^ R package version 4.6.0, random forest models were trained using the ranger^64^ R package version 0.17.0, and logistic regression models were trained using the stats package of R. For conversion of genomic coordinates between hg19 and hg38, the LiftOver^61^ function of the UCSC Genome Web Browser was used.

## Acknowledgements

This work is supported by National Heart, Lung and Blood Institute (NHLBI) Trans-Omics Precision Medicine (TOPMed) Informatics Research Center contract (3R01HL117626-02S1, HHSN268201800002I), National Institutes of Health grants R01HG011031, University of Michigan Training Program in Genome Science T32HG000040. Chan Zuckerberg Institute, and Taubman Institute. We thank NHLBI TOPMed Informatics Resource Center (IRC) and Multi-Omics Working Group for making the whole blood eQTL summary statistics available.

## Competing Interests

The authors declare no competing interests.

## Extended Data Figure Legends

**Extended Data Figure 1.**
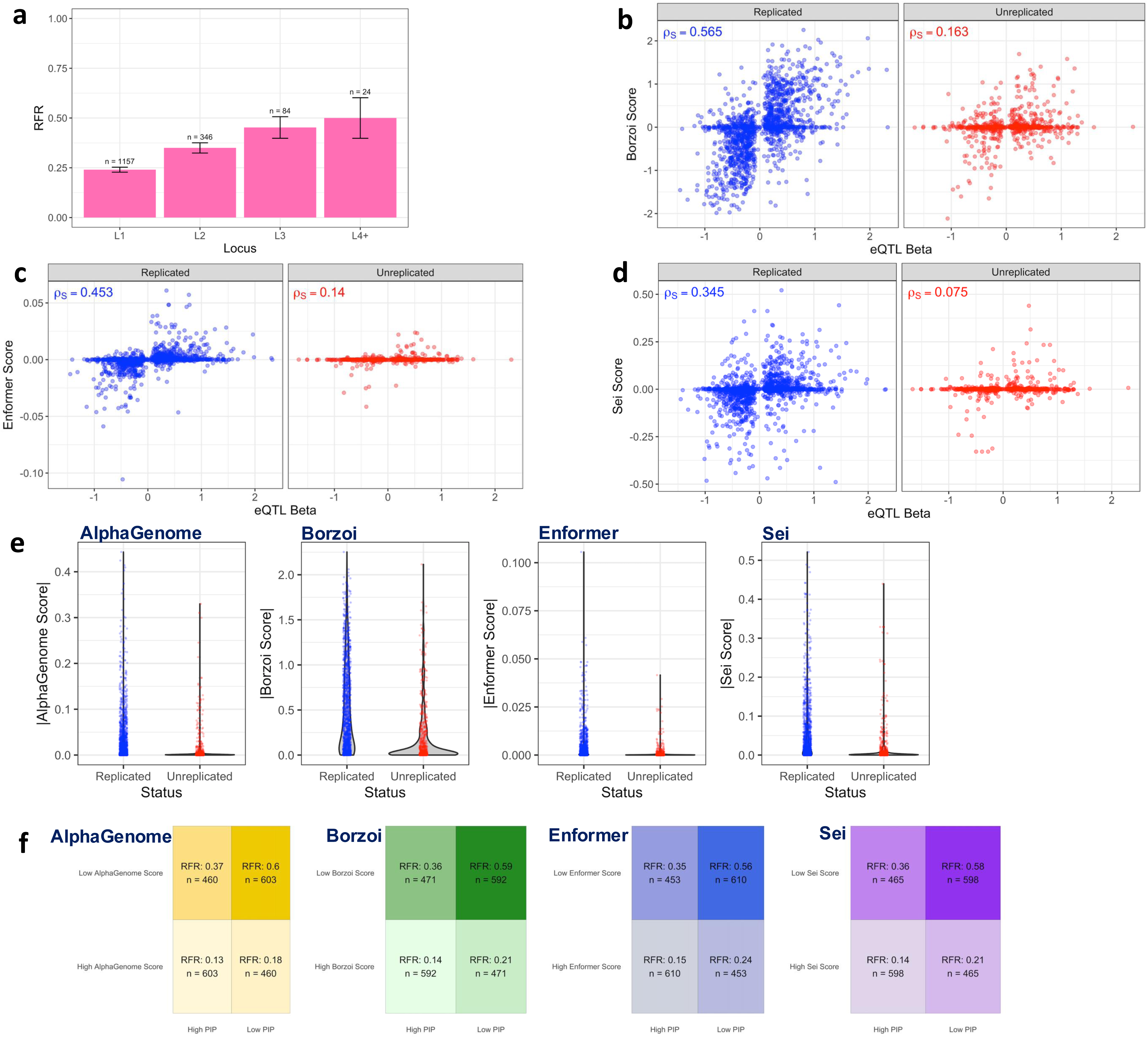
Extended reproducibility analysis high-confidence GTEx blood eQTL credible sets. **(a)** Replication failure rate (RFR) of 1,611 GTEx (v10) blood eQTL credible sets composed of a single SNP with PIP > 99%, stratified by the order of credible sets in SuSiE’s IBSS (Iterative Bayesian Stepwise Selection) algorithm. n indicates the number of credible sets in the category, and the error bars represent estimated standard errors. **(b)** Spearman correlation between eQTL effect sizes and Borzoi scores for 2,126 high-confidence (PIP > 95%) GTEx blood eQTLs, separated by their replication status in TOPMed blood eQTLs. **(c)** Same as (b) but with Enformer scores. **(d)** Same as (b) but with Sei scores. **(e)** The distribution of the magnitude of AlphaGenome, Borzoi, Enformer, and Sei single track scores in 2,126 high-confidence (PIP > 95%) GTEx blood eQTLs, separated by their replication status in TOPMed blood eQTLs. **(f)** Replication failure rate (RFR) of 2,126 high-confidence GTEx (v10) blood eQTL credible sets (PIP > 95%), when credible sets are classified to high/low PIPs and S2O scores based on respective medians, comparing AlphaGenome, Borzoi, Enformer, and Sei.

**Extended Data Figure 2.**
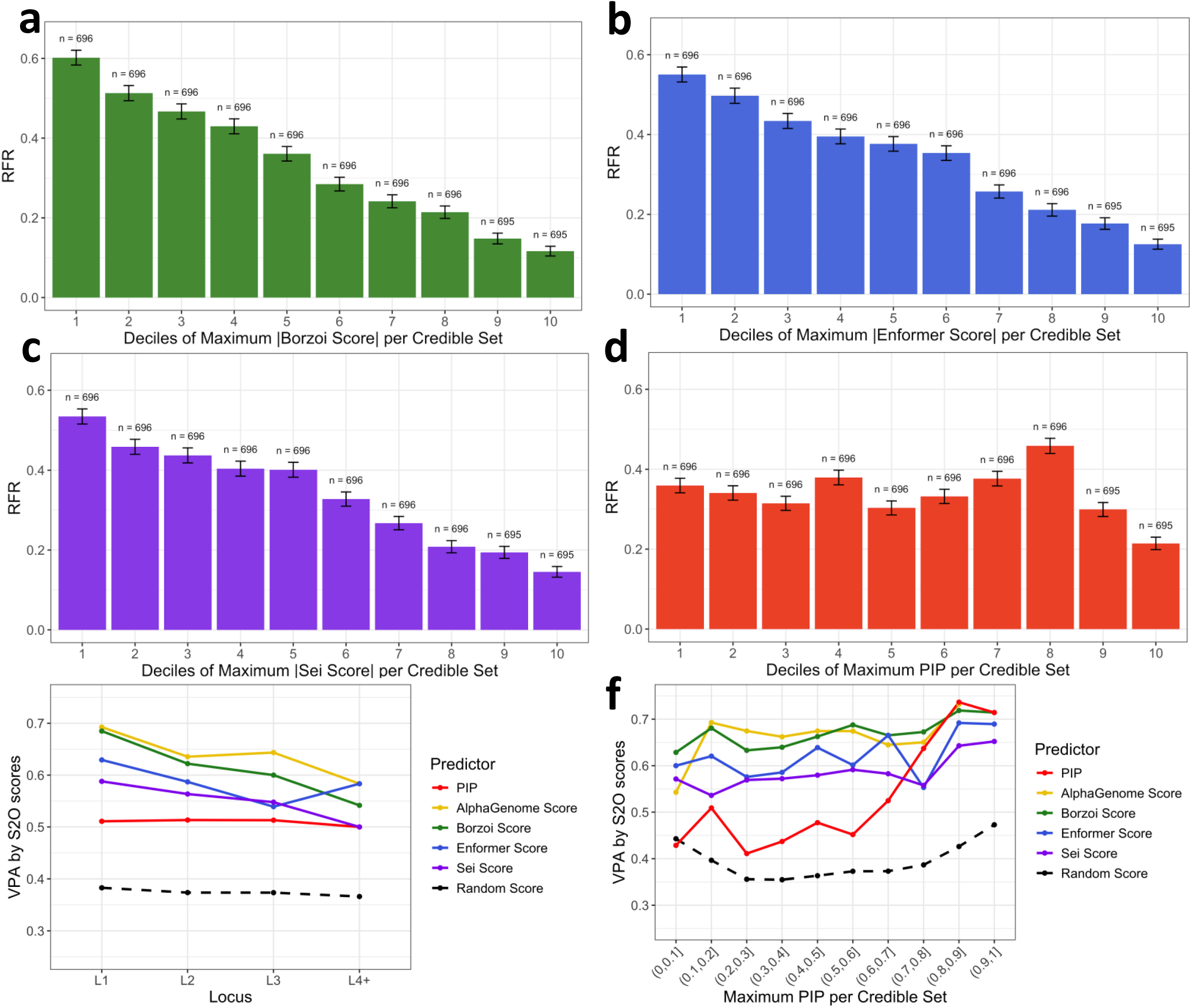
Extended reproducibility analysis of all GTEx blood eQTL credible sets with size ≤ 10. **(a)** Replication failure rate (RFR) of 6,958 GTEx (v10) blood eQTL credible sets with size 10 or less, stratified by the deciles of Borzoi single track scores. **(b)** Same as (a) but with Enformer scores. **(c)** Same as (a) but with Sei scores. **(d)** Same as (a) but with maximum PIP per credible set. **(e)** Variant prioritization accuracy (VPA) of prioritizing replicated variants as the top variant, stratified by the order of credible sets in SuSiE’s IBSS (Iterative Bayesian Stepwise Selection) algorithm, comparing PIP, AlphaGenome, Borzoi, Enformer, and Sei single track scores. Random scores are used as a baseline. **(f)** Same as (d) stratified by the maximum PIP per credible set.

**Extended Data Figure 3.**
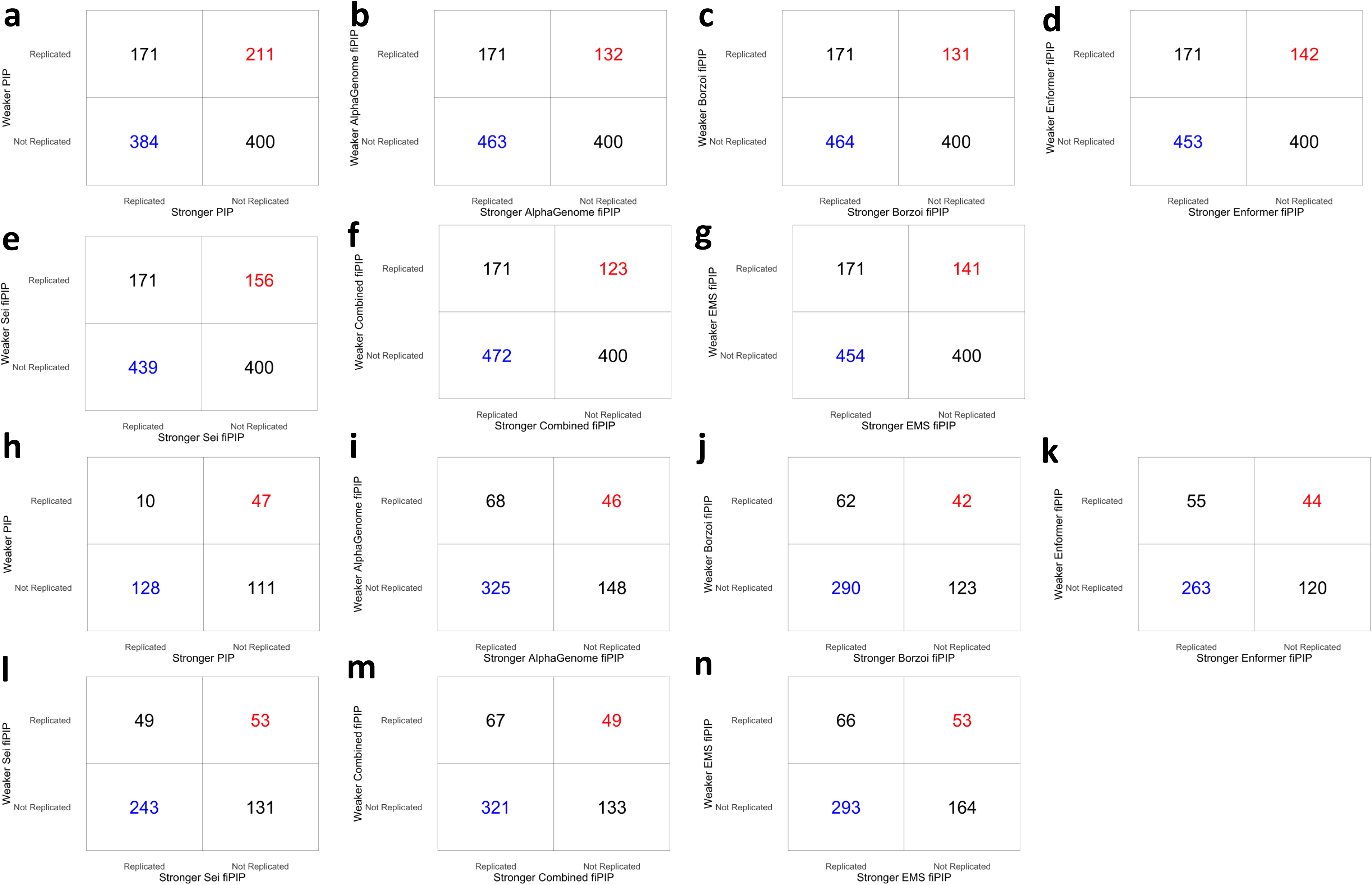
Detailed evaluation of functionally informed fine-mapping for GTEx blood eQTL credible sets with size 2. 2×2 tables of the size 2 credible sets based on the replication status (row) and the relative (fi)PIPs values of the two SNPs in each credible set for **(a)** SuSiE PIP, **(b)** AlphaGenome fiPIP, **(c)** Borzoi fiPIP, **(d)** Enformer fiPIP, **(e)** Sei fiPIP, **(f)** Combined fiPIP, and **(g)** EMS fiPIP. Ties were randomly resolved. Same as (a)-(g) but focusing only on the credible sets containing a SNP with (fi)PIP ≥ 0.9 for **(h)** SuSiE PIP, **(i)** AlphaGenome fiPIP, **(j)** Borzoi fiPIP, **(k)** Enformer fiPIP, **(l)** Sei fiPIP, **(m)** Combined fiPIP, and **(n)** EMS fiPIP.

**Extended Data Figure 4.**
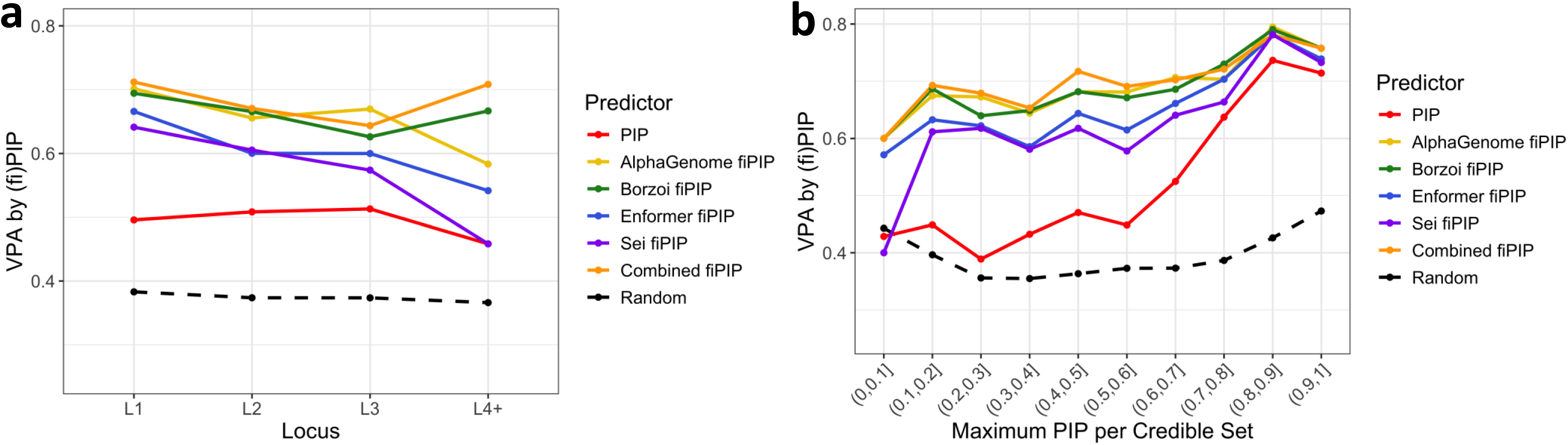
Variant prioritization accuracy (VPA) of prioritizing replicated variants. VPA evaluation using functionally informed posterior inclusion probabilities (fiPIPs) trained by multitrack S2O models, stratified by the **(a)** order of credible sets in SuSiE’s IBSS algorithm, and **(b)** maximum PIP per credible set.

**Extended Data Figure 5.**
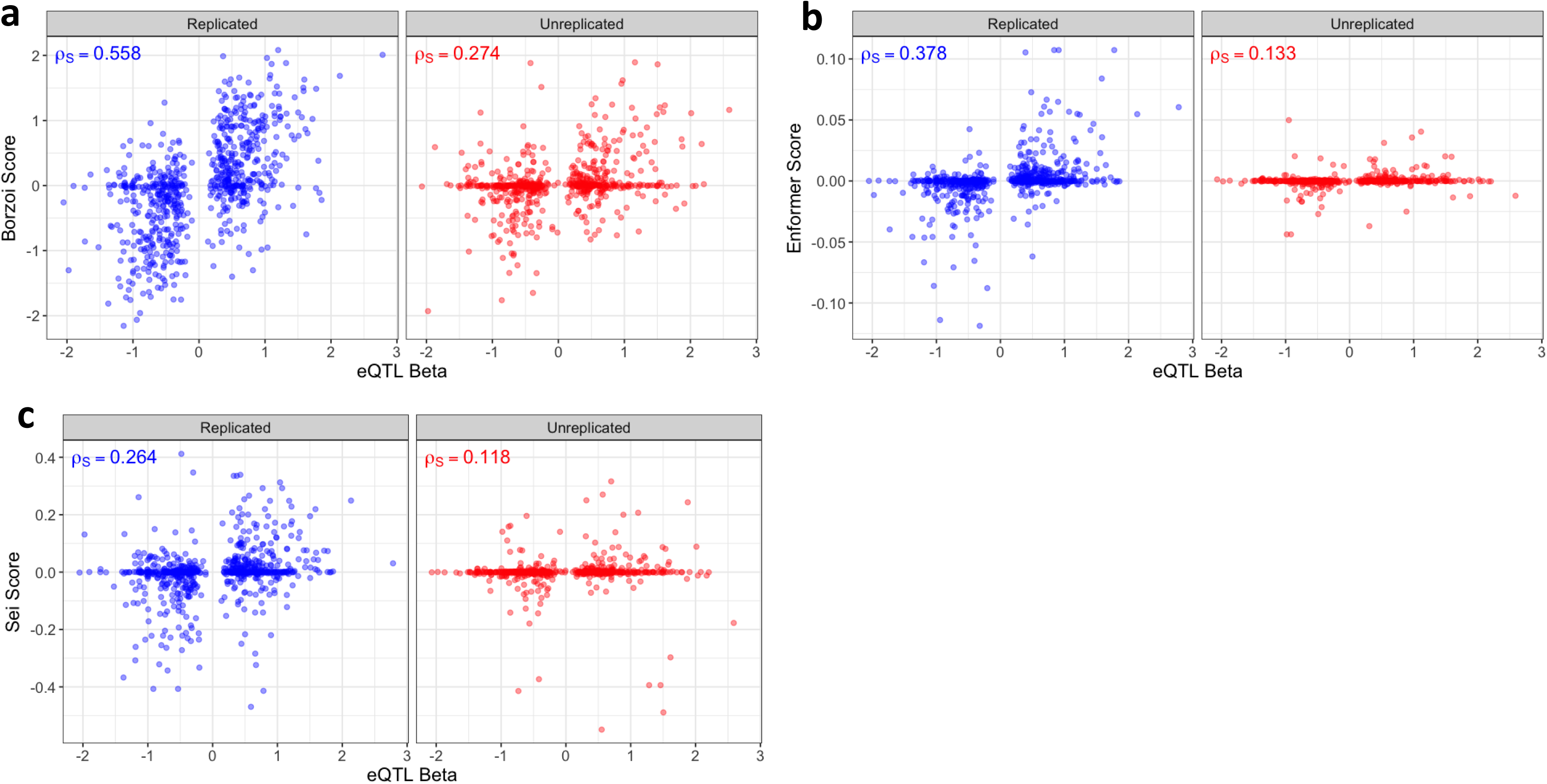
Extended evaluation using GTEx LCL eQTLs. Spearman correlation between eQTL effect sizes and **(a)** Borzoi, **(b)** Enformer, and **(c)** Sei scores for 1,364 high-confidence (PIP > 95%) GTEx LCL eQTLs, separated by their replication status in MAGE LCL eQTLs.

**Extended Data Figure 6.**
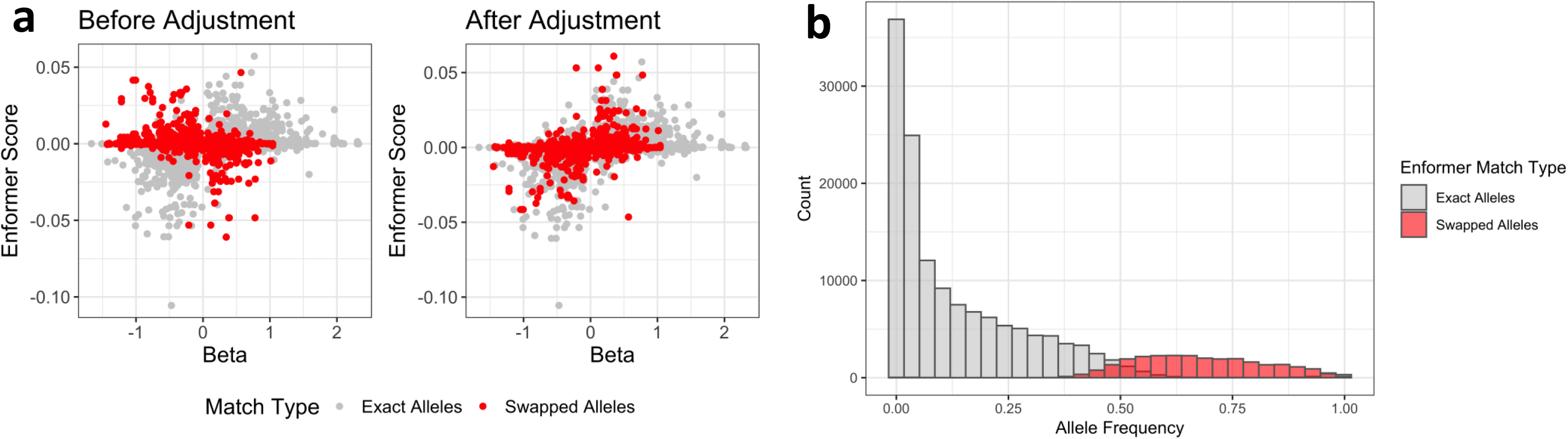
Examination of allele swaps in precomputed Enformer resources. **(a)** Scatterplots between eQTL effect sizes and Enformer single track scores for high-confidence (PIP > 95%) GTEx blood eQTLs, comparing before and after allele swap adjustment based on reference/alternate allele matching. **(b)** Distribution of allele frequencies in GTEx blood eQTLs, separating matched and swapped alleles in precomputed Enformer resources.

## Notes

### Competing Interest Statement

The authors have declared no competing interest.

